# Glycine attenuates impairments of stimulus-evoked gamma oscillation in the ketamine model of schizophrenia

**DOI:** 10.1101/2021.04.15.439976

**Authors:** Moritz Haaf, Stjepan Curic, Saskia Steinmann, Jonas Rauh, Gregor Leicht, Christoph Mulert

**Affiliations:** Department of Psychiatry and Psychotherapy, Psychiatry Neuroimaging Branch (PNB), University Medical Center Hamburg-Eppendorf, Hamburg, Germany; Center of Psychiatry, Justus-Liebig University, Giessen, Germany

## Abstract

Although a substantial number of studies suggests some clinical benefit concerning negative symptoms in schizophrenia through the modulation of NMDA-receptor function, none of these approaches achieved clinical approval. Given the large body of evidence concerning glutamatergic dysfunction in a subgroup of patients, biomarkers to identify those with a relevant clinical benefit through glutamatergic modulation are urgently needed. A similar reduction of the early auditory evoked gamma-band response (aeGBR) as found in schizophrenia patients can be observed in healthy subjects in the ketamine-model, which addresses the putative excitation / inhibition (E/I) imbalance of the diseases. Moreover, this change in gamma-band oscillations can be related to the emergence of negative symptoms. Accordingly, this study investigated whether glycine-related increases of the aeGBR accompany an improvement concerning negative symptoms in the ketamine-model. The impact of subanesthetic ketamine doses and the pretreatment with glycine was examined in twenty-four healthy male participants while performing a cognitively demanding aeGBR paradigm with 64-channel electroencephalography. Negative Symptoms were assessed through the Positive and Negative Syndrome Scale (PANSS). Ketamine alone caused a reduction of the aeGBR amplitude associated with more pronounced negative symptoms compared to placebo. Pretreatment with glycine attenuated both, the ketamine-induced alterations of the aeGBR amplitude and the increased PANSS negative scores in glycine-responders, classified based on relative aeGBR increase. Thus, we propose that the aeGBR represents a possible biomarker for negative symptoms in schizophrenia related to insufficient glutamatergic neurotransmission. This would allow to identify patients with negative symptoms, who might benefit from glutamatergic treatment.

## Introduction

Schizophrenia places a substantial burden on people with this illness, more so as there is no satisfactory pharmacotherapy for some of its core symptoms, namely cognitive and negative symptoms (Correll & Schooler, 2020). A prominent challenge in the pursuit of an effective pharmacotherapy is that patients respond heterogeneously to treatments (Kumar et al., 2020; McCutcheon, Krystal, & Howes, 2020). Therefore, biomarkers to identify the subgroup of patients that would benefit from certain treatments are urgently needed.

A promising approach to new treatments is the modulation of aberrant neural glutamatergic activity in patients with schizophrenia. The glutamate hypothesis of schizophrenia presumes a hypofunction of the N-methyl-d-aspartate receptor (NMDAR) which is essential for glutamate neurotransmission. On this account, there were several attempts to modulate its function both directly (through co-agonists such as glycine or D-serine) and indirectly (through glycine-reuptake inhibitors), which both yielded improvements in negative symptomatology (J. Kantrowitz, 2017; Umbricht et al., 2014). However, none of these pharmacological agents achieved clinical approval despite the repeated confirmation of the viability of this treatment approach through recent studies (Chang, Lin, Liu, Chen, & Lane, 2020; Krogmann et al., 2019). This reinforces the need for biomarkers to detect patients with a putative clinical benefit from glutamatergic modulation.

Gamma band oscillations (GBO) play an essential role in cognition, consciousness, and perception and have been found to be impaired in schizophrenia (Dienel & Lewis, 2019; Uhlhaas & Singer, 2010). The generation of those 30 to 100 Hz frequencies substantially involves NMDARs which are frequently expressed on the inhibitory parvalbumin- (PV+) (Sohal, Zhang, Yizhar, & Deisseroth, 2009) and somatostatin-expressing (SST+) γ-aminobutyric acid (GABA) interneurons (Alherz, Alherz, & Almusawi, 2017). The micro-circuital interplay of the respective interneurons with glutamatergic pyramidal cells (Lisman et al., 2008) then produce said frequency oscillations.

The auditory evoked gamma-band response (aeGBR) is among the sensory evoked GBOs which have been of special interest, given that they do not only reflect sensory processes but appear to be affected by attention and memory (Cho, Konecky, & Carter, 2006; Herrmann, Frund, & Lenz, 2010). The aeGBR appears 25 to 100 ms after an auditory stimulus and exemplifies the top-down operations in sensory processing, as its magnitude is profoundly altered by task difficulty (Mulert et al., 2007). Several studies have proven that aeGBRs are impaired in all stages of schizophrenia (Leicht et al., 2015; Leicht et al., 2010), high risk subjects (Leicht et al., 2016), and even first-degree relatives of people with the illness (Leicht et al., 2011). It is noteworthy, that these detriments are accompanied by a reduced activity of a network including the anterior cingulate cortex (ACC) (Leicht et al., 2015), which has repeatedly been implicated in the pathophysiology of schizophrenia and its cognitive dysfunctions (Reid et al., 2019; Takayanagi et al., 2017).

The ketamine model of schizophrenia offers the opportunity to study this illness without having to test patients. It utilizes the administration of subanesthetic doses of the NMDAR antagonist ketamine to reduce NMDAR dependent glutamatergic neurotransmission, which causes the emergence of schizophrenia-like positive, negative, and cognitive symptoms in healthy volunteers (Krystal et al., 1994) or aggravating symptom severity in patients (Lahti, Koffel, LaPorte, & Tamminga, 1995). This model particularly depicts aberrant GBOs since under physiological conditions ketamine demonstrates the highest affinity for the NMDARs expressing GluN2C and GluN2D subunits, which are most frequently expressed on the aforementioned PV+ and SST+ GABAergic interneurons (Khlestova, Johnson, Krystal, & Lisman, 2016; Kotermanski & Johnson, 2009). The inhibition of the respective interneurons is supposed to result in a disruption of the local microcircuits (Lisman et al., 2008) and leads to an impaired generation of GBO resembling deficiencies observed in patients with schizophrenia (Jadi, Behrens, & Sejnowski, 2016).

Glycine binds to an allosteric binding site of the NMDAR, thus enabling signal transduction following the engagement of glutamate as well as promoting and enhancing the binding of glutamate to the NMDAR (Leeson & Iversen, 1994). These characteristics allow glycine to attenuate neurophysiological impairments in patients suffering from schizophrenia (Greenwood et al., 2018; J. T. Kantrowitz et al., 2018) and neurophysiological impairments related to (induced) NMDAR hypofunction, as animal studies have proven (Lee et al., 2018).

Based on the strong interplay of schizophrenia symptoms, NMDAR dysfunction and receptor co-agonists, this study aimed to delineate the impact of an exogenously induced NMDAR dysfunction on the generation of the aeGBR and the effect of glycine thereon. This was achieved by means of a cognitively demanding auditory choice reaction task after a pretreatment with glycine and during the continuous infusion of ketamine. We hypothesized that glycine-pretreatment mitigates the disturbances of the aeGBR and the interrelated emergence of schizophrenia-like symptoms during ketamine administration in healthy volunteers.

## Results

### Participants

We included 24 healthy male participants in EEG data analysis. Their mean age was 24 years (19 - 32 years, SD 3.7 years) and they had experienced an average of 16.5 educational years (13 - 21 years, SD 2.4 years). All participants were right-handed as assessed by means of the Edinburgh Handedness Scale (mean 77.2 %, 48 - 100 %, SD 16.8 %) and had a mean verbal IQ of 111.5 points (99 - 122 points, SD 6.3 points) according to the German WST-Wortschatztest.

All participants underwent four EEG recording sessions, which differed regarding the pretreatment and continuous infusions. The four corresponding experimental conditions were: (i) placebo-pretreatment followed by placebo (Pla-Pla), (ii) placebo-pretreatment followed by ketamine (Pla-Ket), (iii) glycine-pretreatment followed by placebo (Gly-Pla), and (iv) glycine-pretreatment followed by ketamine (Gly-Ket).

### Behavioral performance

We observed a significant main effect of ketamine on reaction times (F(1,23)=28.4, p<0.001), with increased reaction times occurring during the application of ketamine (Figure 1A). Regarding error rates, a significant main effect of ketamine occurred (F(1,23)=17.7, p<0.001) with ketamine increasing the error rate (Figure 1B). Neither glycine pretreatment nor the interaction between glycine and ketamine affected the behavioral performances.

**Figure 1.**
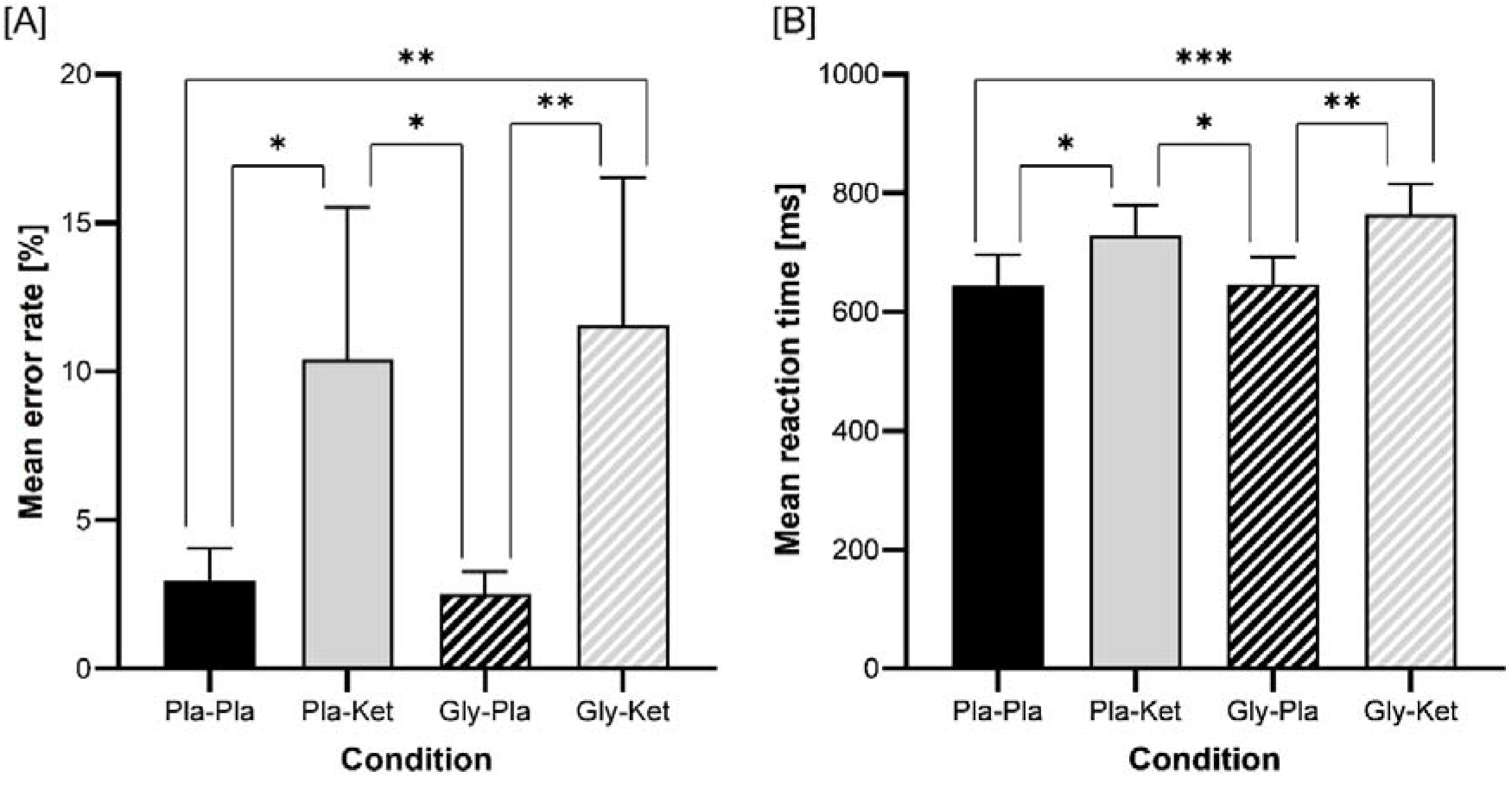
Bar charts of the mean values of the error rate [A] and the reaction time [B]. (***p<0.001, **p<0.01, *p<0.05)

### Ketamine-induced psychopathology

Concerning PANSS total score as well as all factor scores (Figure 2), there was a significant main effect of ketamine (PANSS Total: F(1,23)=121, p<0.001; Positive: F(1,23)=33.7, p<0.001; PANSS Negative F(1,23)=100.9, p<0.001; Disorganization F(1,23)=130.4, p<0.001; Distress F(1,23)=35.5, p<0.001; Excitement F(1,23)=41, p<0.001). Neither glycine pretreatment nor the interaction between glycine and ketamine affected the PANSS total or factor scores.

**Figure 2.**
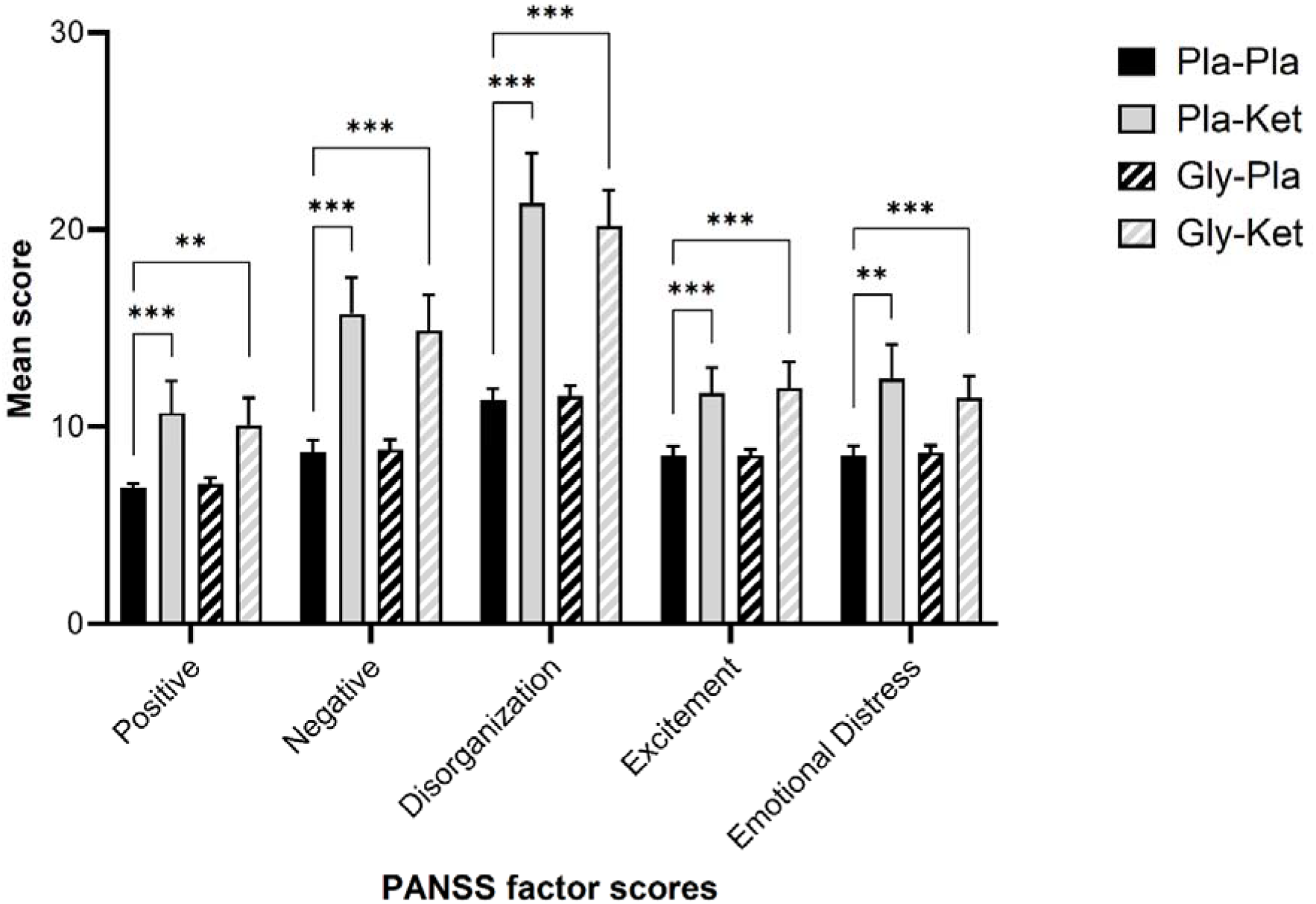
Bar charts of the mean values of the five Positive and Negative Syndrome Scale (PANSS) factor scores. Only significant differences between the Pla-Pla and both ketamine conditions (Pla-Ket and Gly-Ket) are displayed. (***p<0.001, **p<0.01, *p<0.05)

### Auditory evoked GBR amplitude and PLF

Around 50 ms after stimulus presentation in all four conditions, the evoked gamma activity increased at electrode Cz (Figure 3B). Regarding the peaks of the evoked gamma band amplitude, a significant interaction effect between ketamine and glycine occurred (F(1,23)=6.2, p=0.02, η_p_^2^=0.21). Simple main effects analysis revealed a significantly reduced aeGBR amplitude due to the application of ketamine condition following both placebo (p_adjusted_<0.001,95 % CI [-0.203, −0.086]) and glycine (p_adjusted_=0.017 95 % CI [-0.126, −0.014]) pretreatment. Glycine-pretreatment led to an increased aeGBR only when preceding the application of ketamine (p_adjusted_=0.004, 95 % CI [0.022, 0.102]) but not the application of placebo (p_adjusted_=0.718, 95 % CI [-0.058, 0.083]) (Figure 3A).

**Figure 3.**
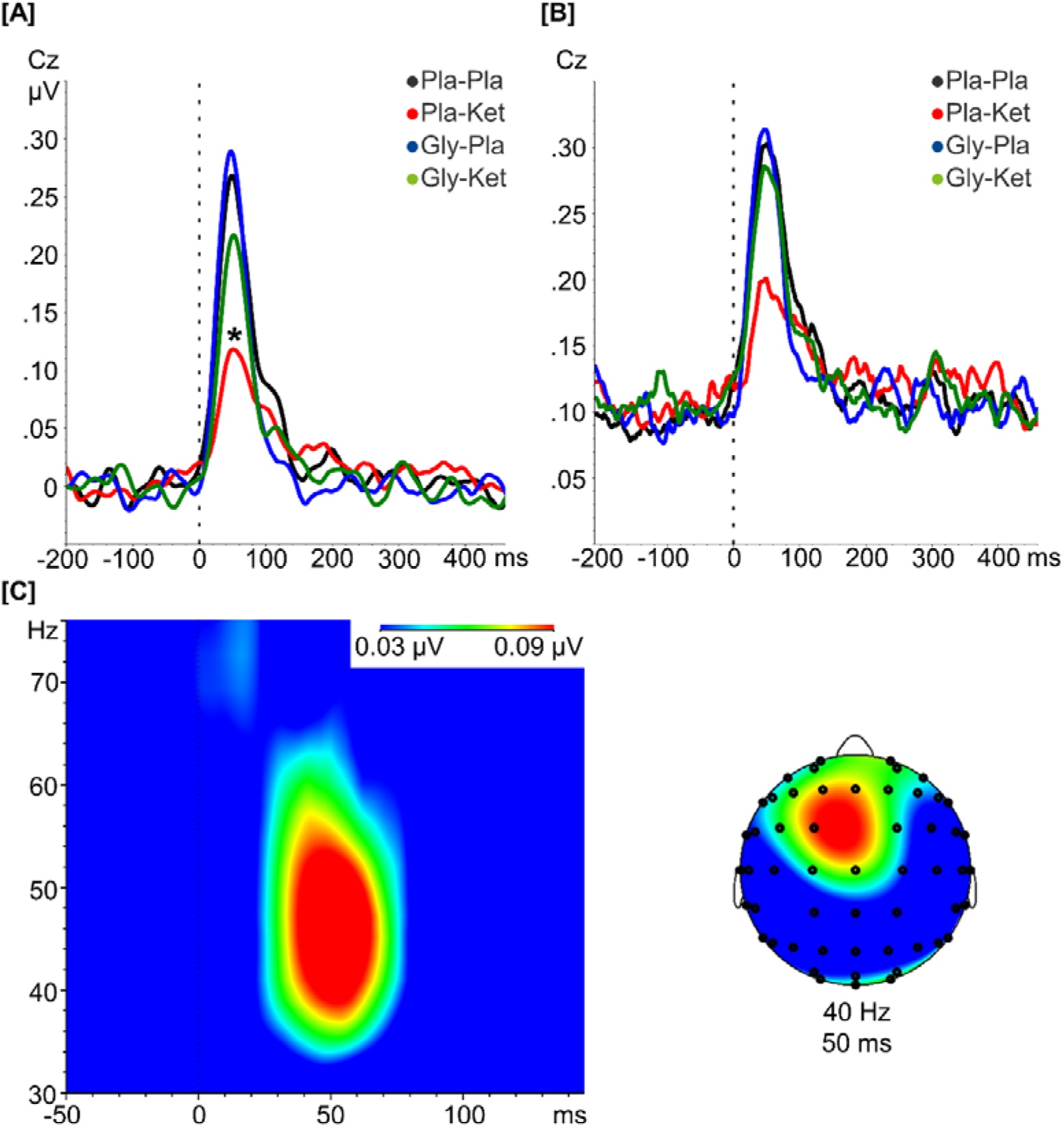
The aeGBR is an increased activity about 50 ms after stimulus presentation (dashed lines). The aeGBR amplitude [A] and phase-locking factor (PLF) [B] are displayed as the results of the wavelet analysis focused on the activity around 40 Hz for the Pla-Pla condition (black line), Pla-Ket (red line), Gly-Pla (blue line) and the Gly-Ket condition (green line). The aeGBR amplitude was significantly altered by an interaction between glycine and ketamine as well as ketamine alone [A] while the PLF was not significantly affected by glycine or ketamine [B]. The time–frequency analysis of the mean difference (Gly-Ket minus Pla-Ket) of the auditory evoked gamma-band response (aeGBR) amplitude and the corresponding topography at 40 Hz and 50 ms after stimulus presentation [C]. (*p<0.05)

Regarding the PLF, neither the interaction between ketamine and glycine (F(1,23)=3.6, p=.071, η_p_^2^=0.14) nor the ketamine main effect (F(1,23)=4.2, p=0.053, η_p_^2^=0.15) nor the glycine main effect (F(1,23)=2.3, p=0.141, η_p_^2^=0.09) reached statistical significance (Figure 3C).

### LORETA whole head analysis

The reduction of the aeGBR due to administration of ketamine (Pla-Pla minus Pla-Ket) involved a significant reduction of gamma activity (30–50 Hz) within the ACC (4 voxels within Brodmann areas 24 and 33, t>3.436, p_adjusted_<0.05, Figure 4A). Glycine-pretreatment before ketamine administration increased the activity of this aeGBR source compared to placebo-pretreatment (Gly-Ket minus Pla-Ket) (Brodmann area 33, t=3.85, p_adjusted_=0.021, Figure 4B).

**Figure 4.**
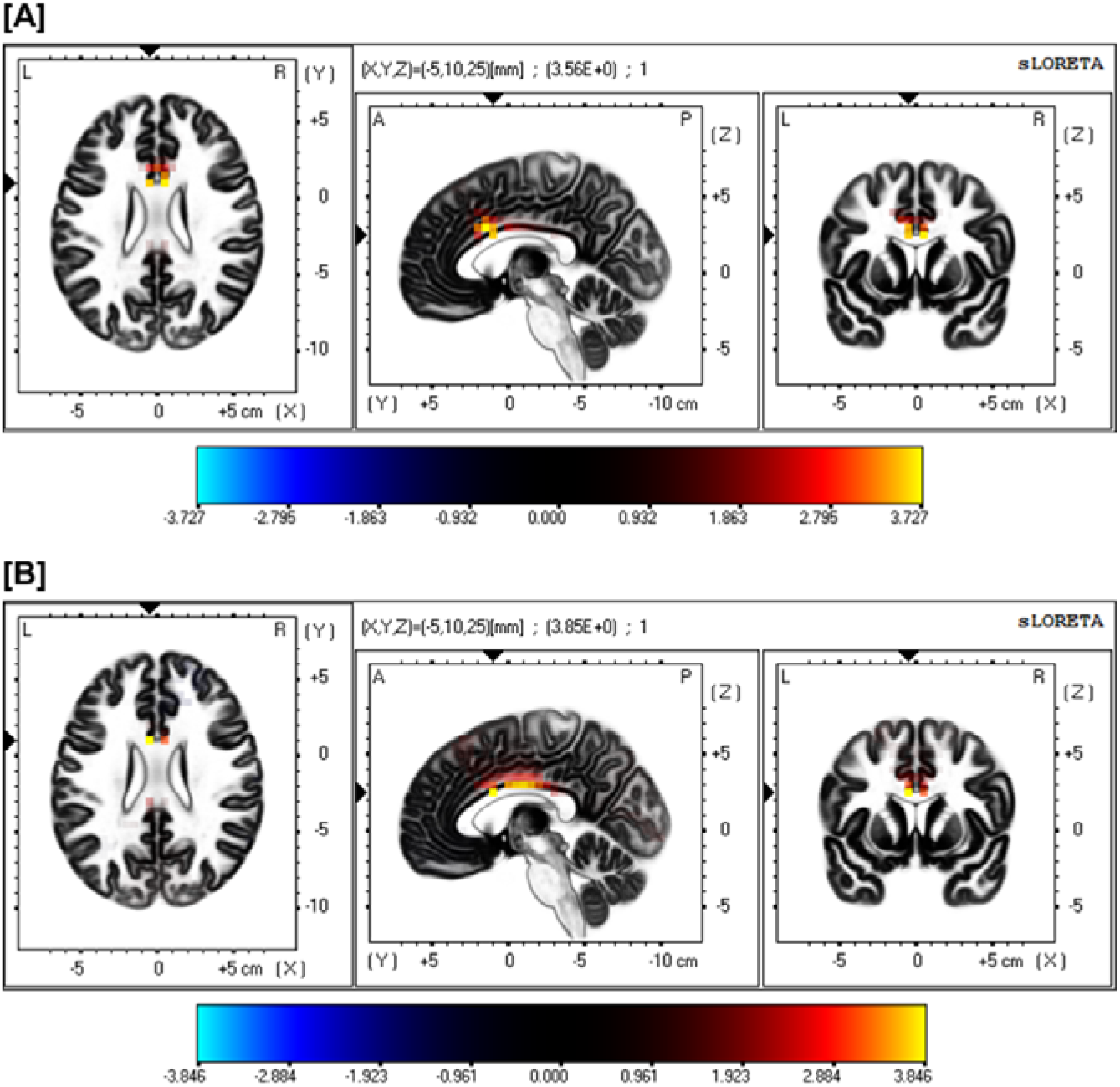
Difference map of low-resolution brain electromagnetic tomography (LORETA) source activity in the gamma-frequency band (30–50 Hz) contrasting the current source density (CSD) of the aeGBR between the Pla-Pla and the Pla-Ket conditions [A] as well as the Gly-Ket and the Pla-Ket conditions [B]. Yellow voxels depict a significantly increased CSD in the anterior cingulate cortex (ACC) comprising the Brodmann areas 24 [A] and 33 [A and B].

### Association between neurophysiological and psychopathological variables

Based on our previous study, we specifically investigated the Pearson correlations between the PANSS negative scores and the relative changes of the aeGBR (Curic et al., 2019). To calculate these relative changes, the Pla-Pla condition was defined as baseline and used as the reference for all contrasts.

Comparing the Pla-Ket and Pla-Pla conditions, increases of the PANSS negative score negatively correlated with the relative reduction of the aeGBR under the influence of ketamine (Pearson’s r=□-0.53, p=0.008) (Figure 5A).

**Figure 5.**
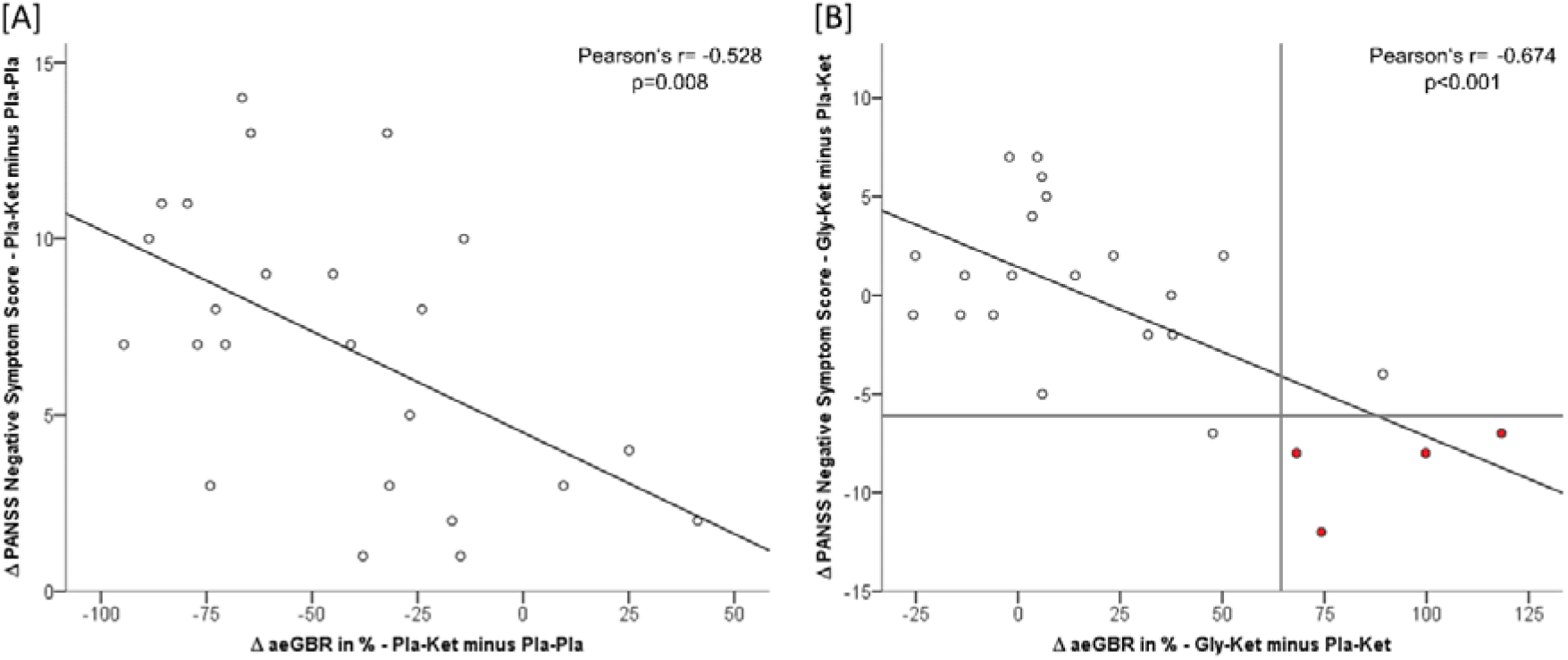
Negative correlation between [A] the relative changes of the aeGBR and changes of the PANSS negative factor comparing the Pla-Ket and the Pla-Pla conditions (Pearson’s r= −0.528 p=0.008) as well as [B] the Gly-Ket and the Pla-Ket conditions (Pearson’s r=□□−0.674 p<0.001). A 64.5 % relative increase of the aeGBR (vertical line) correctly identifies four out of five individuals with a clinically relevant glycine response (red dots) of the PANSS negative factor score (horizontal line, >5.5 points corresponding to the scale intervals of the Clinical Global Impressions-Severity (CGI-S)).

When contrasting the pretreatments of both ketamine sessions (Gly-Ket minus Pla-Ket), the reduction of the PANSS negative score correlated with the relative increase of the aeGBR after pretreatment with glycine (Pearson’s r=□−0.67, p<0.001) (Figure 5B).

In an explorative post-hoc analysis, we divided all individuals into two groups with regard to their glycine-dependent reduction of the PANSS negative score comparing both ketamine sessions, using a clinically significant reduction (of >5.5 points corresponding to the scale intervals of the Clinical Global Impressions-Severity (CGI-S) rating) as the cut-off (Leucht et al., 2019). Employing binary logistic regression, the relative increase of the aeGBR (ΔaeGBR) predicted a clinically relevant attenuation of the PANSS negative score comparing both ketamine conditions (Chi-Square=13.91, df=1, p<0.001). This model determined a ΔaeGBR increase of 64,5 % as the cut-off, which correctly predicted 18 out of 19 (94.7 %) cases where there was no clinically significant attenuation of the PANSS negative score, and four out of five (80 %) cases where there was a significant attenuation.

## Discussion

We have presented evidence that pretreatment with glycine mitigates ketamine-induced impairments of the aeGBR. Furthermore, we have demonstrated that the relative changes of the aeGBR correlate with the severity of schizophrenia-like symptoms associated with the modulation of glutamatergic neurotransmission. While glycine-pretreatment did not affect behavioral performance and PANSS factor scores at the group level, the aeGBR amplitude allowed us to identify individuals with a relevant psychopathological benefit from the pretreatment with glycine.

Cortical and subcortical GBO both at rest or task-driven (e.g. aeGBR) are critically involved in cognitive functions such as working memory (Howard et al., 2003; van Vugt, Schulze-Bonhage, Litt, Brandt, & Kahana, 2010) and are found to be altered in schizophrenia (Dienel & Lewis, 2019; Uhlhaas & Singer, 2010). The early aeGBR is known to be reduced across all stages of schizophrenia (Leicht et al., 2015; Leicht et al., 2010) and corresponding impairments can be found in the ketamine model (Curic et al., 2019). Moreover, the activity of the aeGBR generators within the dACC is reduced in patients with schizophrenia (Leicht et al., 2015; Leicht et al., 2010) and in healthy subjects following the acute infusion of ketamine (Curic et al., 2019). Accordingly, we were able to replicate these findings in the present study.

Based on these observations the aeGBR has been proposed as a correlate to disrupted glutamatergic neurotransmission in schizophrenia, since the generation of oscillations in the gamma frequency range depends on a feedback-loop encompassing pyramidal cells and GABAergic interneurons (Lisman et al., 2008). This premise is further supported by our finding that the alleviation of ketamine-related aeGBR impairments follows the modulation of glutamatergic neurotransmission through the NMDAR co-agonist glycine. Yet, the effect of glycine on humans had neither been studied in the ketamine model of schizophrenia (Haaf, Leicht, Curic, & Mulert, 2018) nor in light of aberrant GBO. Regarding another neurophysiological measure, the acute administering of glycine normalized the reduced duration mismatch negativity (MMN) amplitudes in patients suffering from schizophrenia (Greenwood et al., 2018). This was paralleled by findings of an improved frequency MMN after continuous treatment with D-serine (another NMDAR glycine-binding site agonist) (J. T. Kantrowitz et al., 2018).

In the aforementioned feedback-loop, PV^+^ and SST^+^ interneurons significantly contribute to the model of NMDAR-dysfunction in schizophrenia, since these subtypes express high densities of NMDARs containing either GluN2C (PV^+^) or GluN2D (SST^+^) subunits for both of which ketamine demonstrates a higher affinity under physiological conditions compared to other subunits (Bygrave, Kilonzo, Kullmann, Bannerman, & Kätzel, 2019). Conversely, both types of interneurons are reduced in schizophrenia according to postmortem studies (Konradi et al., 2011). One might speculate that the disruption of inhibitory feedback might lead to an excessive yet uncoordinated downstream release of glutamate through pyramidal cells, causing an imbalance of glutamatergic excitation and GABAergic inhibition (E/I imbalance) (Kehrer, Maziashvili, Dugladze, & Gloveli, 2008). This offers a feasible explanation for the counterintuitive observation that the acute administering of ketamine increases extracellular glutamate levels in several brain regions assessed by means of 1H-MRS (Stone et al., 2012), paralleled by findings in schizophrenia (Merritt, Egerton, Kempton, Taylor, & McGuire, 2016). Further, the reports of increased resting-state gamma band oscillations in schizophrenia (Andreou et al., 2015; Baradits et al., 2019) as well as corresponding findings in the ketamine model (Bianciardi & Uhlhaas, 2021) – seemingly contradicting the notion of a hypo-glutamatergic state – might be attributed to a putative lack of synchronized neuronal activity. Hence, the concurrent inhibition of PV^+^ and SST^+^ GABAergic interneurons offers an explanation implementing both the incapacity to increase GBO following a cognitive-demanding stimulus and elevated GBO at rest in light of the E/I imbalance resembling a reduced signal/noise ratio.

Accordingly, animal studies found the ability to increase stimulus-evoked gamma to be diminished ensuing global disruption of NMDAR signaling or the administering of NMDAR-antagonists or selective reduction of PV^+^-cell activity by means of either optogenetic methods (Sohal et al., 2009) or gene knockout (Bygrave et al., 2019). Whereas, the results of animal studies focusing on SST^+^ interneurons remain limited and inconclusive (Alherz et al., 2017). Nonetheless, a dysfunction restricted to a specific neuron subtype is likely not enough to explain the pathophysiology and complex neurophysiological changes seen in schizophrenia (Bygrave et al., 2019).

Bearing all considerations outlined above in mind, the association between an impaired early aeGBR and the emergence of schizophrenia-like negative symptoms as seen in this study leads to the presumption that glutamatergic dysfunction, aberrant neural oscillations, and negative symptoms are interdependent. Accordingly, our results contribute to a growing body of evidence that suggests an interrelation of impaired evoked GBO in a cognitive demanding task based on NMDAR dysfunction and negative symptoms as reported in both the ketamine model (Curic et al., 2019) and schizophrenia itself (Leicht et al., 2015). These assumptions are further substantiated by our discovery that the NMDAR co-agonist glycine mitigates ketamine-related aeGBR impairments which in turn presents as a recovery of PANSS negative scores.

Remarkably, four out of five individuals showing a clinically relevant attenuation of the PANSS negative score (Leucht et al., 2019) following glycine-pretreatment could be identified post-hoc by the relative aeGBR-increase with a high specificity and sensitivity (Figure 5B). Hence, the aeGBR could serve as a putative biomarker for target-engagement of glutamatergic remedies in schizophrenia and could help to identify individuals who might benefit from the corresponding treatment. Insufficient target-engagement or competing target-engagement of co-administered medication (i.e. clozapine) might explain the varying results of glycine and D-serine treatments in schizophrenia (J. Kantrowitz, 2017). Nevertheless, further verification of our observations is required given the heterogenous response to the paradigm and considering we are the first to report a mitigating effect of a glutamatergic substance on psychopathological and neurophysiological aberrations in the ketamine model of schizophrenia.

The ketamine-induced increases in reaction time and error rates accompanied by the occurrence of schizophrenia-like symptoms, assessed by means of the PANSS, add to an extensive portfolio of data corroborating the presumptive overlap of schizophrenia itself and the related ketamine model (Frohlich & Van Horn, 2014). Glycine-pretreatment however, paralleled by unsuccessful clinical trials of glutamatergic agents (J. Kantrowitz, 2017), did not reverse these effects at the group level, while clearly mitigating ketamine-associated aeGBR reductions. Nonetheless, glycine-pretreatment reduced PANSS negative scores in the subset of individuals that also demonstrated a profound increase of the aeGBR, whilst no individuals with an increase of the PANSS negative factor by more than two points responded to glycine-pretreatment regarding the aeGBR.

Finally, our study was limited by several factors that merit consideration. First, the small sample size of the experiment represents an important limitation, possibly contributing to the fact that we did not observe a mitigating effect of glycine on behavioral results (e.g., error rate or reaction time) due to medium or small effect sizes. Moreover, we acknowledge that the results of our post-hoc binary logistic regression analysis demand replication with a larger sample size. Secondly, the psychotomimetic effects of ketamine could have led to the partial unblinding of participants. While blinding for the pretreatment was thoroughly realized, further studies could implement dopaminergic agonists or non-glutamatergic psychotomimetic agents in place of saline infusions as the control for ketamine.

In conclusion, this is the first study to report a normalizing effect of glycine, an NMDAR co-agonist, on ketamine-related decreases of the aeGBR peak amplitude and on the activity of its generator located in the ACC in healthy human subjects during the performance of a cognitively demanding auditory choice reaction task. Further, we found an association between glycine-related aeGBR increases and improvements in schizophrenia-like negative symptoms. Remarkably, an increase of the aeGBR predicted a clinically relevant psychopathological response to glycine-pretreatment. This points to the applicability of the aeGBR as a putative rapid biomarker for responders to a glutamatergic therapy for negative symptoms. To this end, our intriguing results call for the investigation of the transferability of this effect to patients suffering from schizophrenia.

## Methods and Materials

### Participants

Twenty-six healthy male participants were enrolled in this study. The general procedure was approved by the Ethics Committee of the Medical Association Hamburg and carried out in accordance with the latest version of the Declaration of Helsinki. Written informed consent was obtained from all participants after the nature of the procedures had been fully explained. One participant dropped out due to adverse effects caused by ketamine (dissociative effect/headache). Another participant had to be excluded due to poor EEG data quality. Volunteers were either students or medical staff of the University Medical Center Hamburg and received a monetary compensation for the EEG recording sessions.

The inclusion criteria included male gender, an age between 18 – 40 years, right-handedness, normal or corrected visual acuity and German at native speaker level.

Exclusion criteria were any acute or previous psychiatric disorders, a family history of schizophrenia or bipolar disorder, the use of illegal drugs, active medication or health conditions representing a contraindication to the administration of ketamine.

The required sample size was calculated using G*Power 3.1 (Faul, Erdfelder, Lang, & Buchner, 2007). With respect to the reduction of the aeGBR amplitude through the application of ketamine and the potential attenuation of this effect via pretreatment with glycine, we expected a medium effect size in the range of η_p_^2^=0.1. This estimation was based on a previous study comparing the effect of the administration of ketamine on the aeGBR to placebo (d=0.6) (Curic et al., 2019). We planned to apply regular statistical analyses and error probabilities (ANOVA, α=0.05; 1-β=0.95). Thus, the sample size analyses led to a total sample size of n = 21 required for a repeated measure, within factors ANOVA for one group and four measurements. To completely counterbalance the order of the four experimental conditions, we decided to include 24 participants.

### Study Design

This study followed a double-blind (regarding pretreatment), randomized, placebo-controlled crossover design. All participants underwent four EEG recording sessions, which differed regarding the pretreatment and continuous infusions. The four corresponding experimental conditions were: (i) placebo-pretreatment followed by placebo (Pla-Pla), (ii) placebo-pretreatment followed by ketamine (Pla-Ket), (iii) glycine-pretreatment followed by placebo (Gly-Pla), and (iv) glycine-pretreatment followed by ketamine (Gly-Ket). The order of sessions was randomized and overall counterbalanced. The glycine-pretreatment was administered at a dosage of 200 mg/kg bodyweight (Greenwood et al., 2018) as an intravenous infusion in 500 ml 0.9 % sodium chloride (NaCl) solution over one hour. Placebo was administered analogously as a NaCl infusion. Both pretreatments were prepared by an unblinded third person prior to the recording sessions and the ready-to-use infusions were indistinguishable. Subsequently, during the ketamine sessions a subanesthetic dose of S-ketamine hydrochloride (Ketanest^®^ S, Pfizer) was administered intravenously in a 0.9 % NaCl solution for a duration of 75 minutes. The infusion was started with an initial bolus of 10 mg over 5 minutes followed by a maintenance infusion of 0.006 mg/kg/min, reduced by 10 % every 10 minutes (Curic et al., 2019). Placebo was administered analogously as NaCl infusions. Heart rate, blood pressure and oxygen saturation were continuously monitored during all sessions. The clinical raters were blinded with respect to the pretreatment but not regarding the continuous infusion condition due to the clinical effects of ketamine.

### Psychometric assessment

The psychiatric symptomatology was assessed by an experienced psychiatrist using the Positive and Negative Syndrome Scale (PANSS) (Kay, Fiszbein, & Opler, 1987) after each recording session. Based on our previous studies the PANSS scores were evaluated using the five-factor model (van der Gaag et al., 2006).

### Stimuli

An attentionally demanding auditory reaction task used in previous studies of our group was employed (Curic et al., 2019; Leicht et al., 2015; Mulert et al., 2007). Three tones of different pitch (800, 1000 and 1200 Hz; 40 repetitions of each) were generated and presented to participants, who were instructed to respond as quickly and as accurately as possible to the low or the high tone (target tones) by pressing a corresponding button. Reaction times and errors were registered and only trials with correct responses to target tones were considered for further analyses.

### EEG recording

In a sound-attenuated and electrically shielded room subjects were seated in a reclined chair with a head rest with their eyes open and asked to look at a fixation cross presented at a monitor 1 m in front of them. The EEG was recorded at a sampling rate of 1000 Hz including an analog band-pass filter (0.1 – 1000 Hz), with 64 active electrodes mounted on an elastic cap (actiCap, Brain Products, Gilching, Germany) in an extended 10/20 system (Curic et al., 2019), using the Brain Vision Recorder software Version 1.21 (Brain Products). Eye movements were recorded through four EOG channels. An electrode at the FCz position was used as reference, the electrode at position AFz served as ground. Impedances were kept below 5 kΩ.

### Data Analysis

Data preprocessing and analysis was carried out using the Brain Vision Analyzer (BVA) Version 2.1 (Brain Products). The preprocessing procedure was conducted in accordance with previous studies (Curic et al., 2019; Leicht et al., 2010). After band-pass filtering (1–80 Hz) and down-sampling to 500 Hz, a topographic interpolation (spherical splines) of up to five channels was performed (the mean number of interpolated channels did not differ between conditions). The channels were selected for interpolation in accordance with the procedure mentioned above. An independent component analysis (ICA) was performed to identify and remove blinks, drifts, muscle artifacts and saccadic spike potentials based on their characteristic topographies, time courses, and frequency distributions (the mean number of removed ICA components did not differ between conditions). The continuous EEG was segmented into epochs of 3000 ms starting 1000 ms prior to the auditory stimulus. Segments including amplitudes exceeding ±70 μV within a 600 ms window starting 200 ms pre-stimulus in any channel were automatically rejected. After re-referencing to common average reference and baseline correction (using an interval of −210 to −10 ms pre-stimulus), averaged event-related potential (ERP) waveforms were computed. Only waveforms based on at least 35 segments were accepted and included in further analyses.

### aeGBR amplitude and phase-locking factor (PLF)

The aeGBR amplitude and PLF were computed using a complex Morlet wavelet transformation (Morlet parameter c=5, instantaneous amplitude (Gabor) normalization) as applied in several previous studies (Curic et al., 2019; Leicht et al., 2010; Mulert et al., 2010). This wavelet transformation was performed on averaged ERPs to obtain the phase-locked evoked gamma amplitude. The analysis followed previously published and well-established procedures and extracts the highest value within the timeframe of 30–100 ms post-stimulus at the electrode Cz of the wavelet layer with the central frequency of 40 Hz (frequency range 32–48 Hz) (Curic et al., 2019; Leicht et al., 2015; Leicht et al., 2011; Leicht et al., 2010; Leicht et al., 2016; Mulert et al., 2007). The PLF was calculated by performing a wavelet transformation at the single trial level and extracting complex-phase information with all vector lengths normalized to the unit circle before averaging the phase information (Curic et al., 2019; Leicht et al., 2010). Gamma PLF peaks were defined as the highest value of the wavelet layer centered around 40 Hz within the timeframe of 30–100 ms post-stimulus at the electrode Cz.

### Source analysis of the aeGBR

Source-space localization analyses were performed with the low-resolution brain electromagnetic tomography (LORETA) KEY software package v2017 (Pascual-Marqui, 2002). The EEG source localization of the aeGBR across 30-50 Hz was executed for every subject within a time window of 30 to 70 ms post-stimulus.

The voxel-wise comparison of cortical activities between conditions was conducted using a one-tailed *t*-test for paired groups (statistical significance threshold *p*□<□0.05) provided in the sLORETA software. A statistical nonparametric mapping randomization method was used to automatically adjust for multiple comparisons with a Fisher’s random permutation test with 5000 randomizations.

### Statistical analyses

All statistical analyses were performed using IBM SPSS Statistics Version 24. Comparisons of reaction times and error rates, PANSS scores, aeGBR amplitude, and PLF between all conditions were conducted using 2 (ketamine) x 2 (glycine) repeated measure analyses of variance (RM-ANOVA). All follow-up simple main effect analyses and post-hoc contrasts were subject to a Bonferroni correction.

## Acknowledgements

This work was prepared as part of Moritz Haaf’s dissertation at the University of Hamburg.

